# Structure of the hepatitis C virus E1/E2 envelope proteins in a homodimeric complex

**DOI:** 10.1101/2023.12.27.573427

**Authors:** Elias Honerød Augestad, Christina Holmboe Olesen, Christina Grønberg, Andreas Soerensen, Rodrigo Velázquez-Moctezuma, Margherita Fanalista, Jens Bukh, Kaituo Wang, Pontus Gourdon, Jannick Prentoe

**Affiliations:** Copenhagen Hepatitis C Program (CO-HEP), Department of Infectious Diseases, Copenhagen University Hospital, Hvidovre, Denmark; Copenhagen Hepatitis C Program (CO-HEP), Department of Immunology and Microbiology, Faculty of Health and Medical Sciences, University of Copenhagen, Copenhagen N, Denmark; Department of Biomedical Sciences, Faculty of Health and Medical Sciences, University of Copenhagen, Copenhagen, Denmark; Department of Experimental Medical Science, Lund University, Lund, Sweden

## Abstract

Worldwide, 58 million individuals suffer from chronic hepatitis C virus (HCV) infection, a primary driver of liver cancer. The HCV envelope proteins, E1 and E2, form a heterodimer, which is the target for neutralizing antibodies. Despite high-resolution structural models of partial heterodimer elements, the structural landscape of higher-order E1/E2 oligomers remains unexplored. We determined a ~3.5 Å cryo-electron microscopy structure of membrane-extracted HCV E1/E2 in a homodimeric arrangement. This structure includes detailed information on the homodimer interface, the E2-binding pocket for hypervariable region 1, antigenic site 412 conformation, and the organization of the E1/E2 transmembrane regions, including one internal to E1. This higher-order E1/E2 assembly could play a pivotal role in the design of novel vaccine antigens better mimicking E1/E2 complexes on the HCV particle.

## Main Text

HCV is an enveloped positive-stranded RNA virus that belongs to the *Flaviviridae* family, classified into 6 clinically important major variants, with genotypes 1 and 3 accounting for most infections, worldwide *(1)*. The viral genome encodes a polyprotein that undergoes processing to yield ten proteins, including the two envelope glycoproteins of the virion, E1 and E2, which form non-covalent heterodimers. E1/E2 complexes facilitate viral entry and serve as the primary targets for neutralizing antibodies (NAbs). During the entry process, E2 has been shown to engage the HCV entry co-receptors, tetraspanin CD81 and scavenger receptor class B, type I, facilitating translocation of the virion to the cell tight junction *(2)*. It remains unclear whether E1 or E2 is the protein responsible for virus fusion with the target cell *(3, 4)*. Conformational dynamics of E1/E2 complexes are likely affecting HCV NAb epitope accessibility, but the structural basis for this is missing *(5)*.

Although the development of an effective vaccine against HCV has proven challenging, it is apparent that worldwide control will require implementation of vaccine strategies protecting from chronicity *(6)*. A major hindrance of developing vaccine candidates inducing effective Nabs is the limited understanding of native E1/E2 protein topology, and, in particular, that higher-order oligomeric states of E1/E2 remain undefined. While structural information exists for parts of monomeric E1 and E2, as well as most of the E1/E2 heterodimer *(4, 7-15)*, vaccine antigens based solely on these incomplete structures may not represent ideal starting points for rational vaccine design. This is because relevant NAb epitopes may be missing, while epitopes that are buried in native oligomeric E1/E2 complexes are exposed, leading to the elicitation of low-potency or non-neutralizing NAbs. Given the correlation between spontaneous HCV clearance and the presence of broadly reactive NAbs (bNAbs) *(16, 17)*, directing the immune response towards conserved epitopes of native E1/E2 complexes may be key in the development of an effective vaccine *(18)*.

The assembly of higher-order complexes of E1/E2 heterodimers has long been debated. Recently, the structure of HCV E1/E2 derived from the genotype 1a isolate, AMS0232, was solved, shedding light on hydrophobic and conserved regions within the ectodomain that are exposed and likely involved in oligomerization *(19)*. It has been speculated that E1/E2 trimerizes through the transmembrane (TM) domain of E1, but the topology of complexes formed by the E1/E2 heterodimer remains elusive *(20, 21)*. Furthermore, the position of the N-terminal part of E2 is unknown. This part is comprised of two distinct segments: a sequence-diverse region called hypervariable region 1 (HVR1, residues 384-410), followed by a short, sequence-conserved, but very flexible segment known as antigenic site 412 (AS412, residues 412-423) (Fig. 1A). These regions play critically linked, but incompletely understood, roles in HCV NAb evasion, and serve as regulators of early HCV co-receptor interactions *(22)*. It has been proposed that these segments are intrinsically disordered *(23)*, as they display flexibility in transmembrane (TM)-truncated, ectopically expressed E2 *(8)*. However, our understanding of their characteristics within higher-order E1/E2 complexes remains limited.

**Fig. 1.**
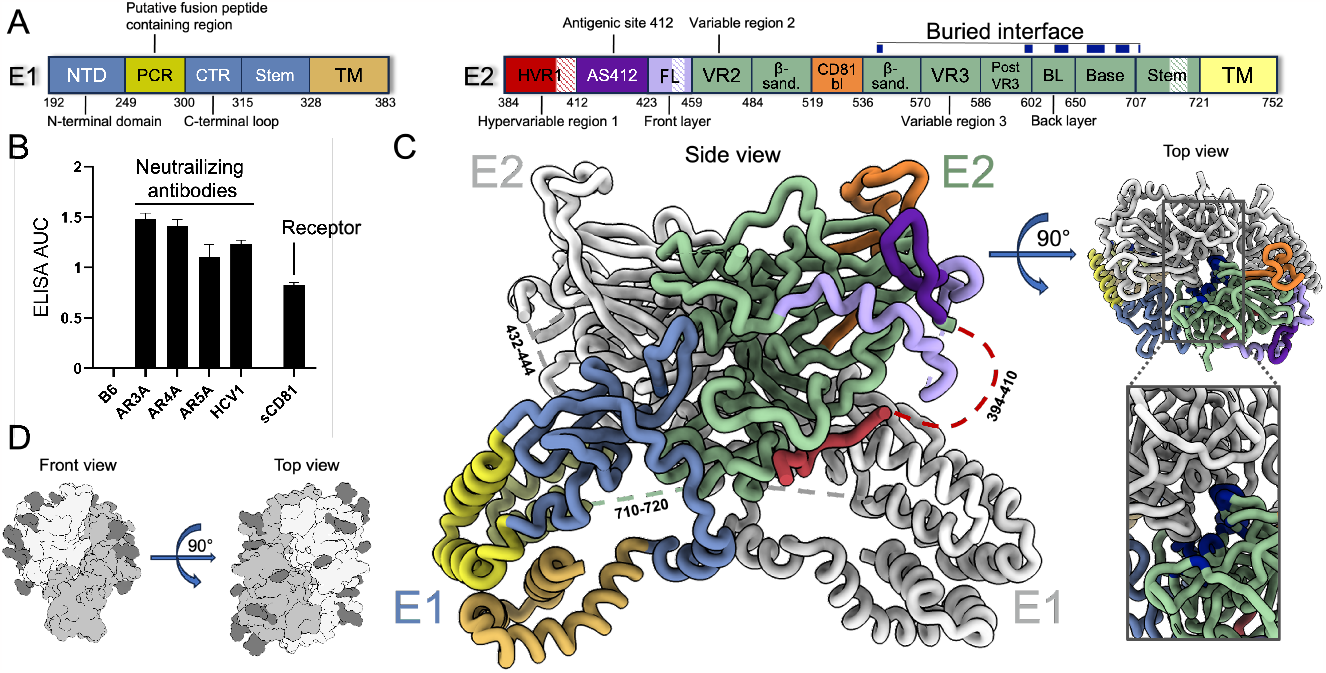
Cryo-EM structure of HCV E1/E2 heterodimers in a homodimeric complex. (**A**) Schematic representation of E1 and E2. The E1 and E2 subunits are shown in blue and green, respectively, with the regions for which this model provides novel structural information highlighted in different colors and missing regions in white with colored stripes. The blue boxes on top of the E2 schematic represent the part of the E2 regions buried within the interface. The same color coding is used in (C). (**B**) Binding of NAbs and sCD81 measured by ELISA expressed as area-under-the-curve (AUC). (**C**) Model of E1/E2 in a homodimeric complex. The missing regions (residues 394-410, 432-444 and 710-720) are depicted in broken lines. (**D**) Display of glycan distribution on the E1/E2 homodimeric complex. One E1/E2 heterodimer is represented in light gray and the other in gray, with the glycan density depicted in dark gray.

In this study, we purified and determined the cryo-electron microscopy (cryo-EM) structure of E1/E2 in a homodimeric complex. This provides detailed information of an anti-parallel arrangement of two E1/E2 heterodimers, with HVR1 extending into a cavity on E2 with AS412 locked in a β-hairpin conformation. This complex offers several key insights into the biology of HCV and provides a framework for designing vaccine antigens focused on targeting epitopes exposed on HCV particles.

## Results

### Purification and structural determination of HCV E1/E2 in a homodimeric complex

We produced HCV envelope proteins in HEK293T cells and ran detergent-extracted HCV E1/E2 from different HCV isolates on blue native (BN) PAGE, visualizing the protein with the conformation-dependent NAbs, AR3A *(24)* for E2, as well as AR4A *(25)* for E1/E2. E1/E2 from the genotype 1a isolate, H77 *(26)*, gave a distinct band of 300-400 kDa (fig. S1A). The genotype 3a isolate, S52 *(27)*, gave a similar but more diffuse band that became more defined by the replacement of residues from the H77 C-terminal TM helices of E1 and E2 (S52mod, fig. S1B). The E1/E2 complex was destabilized by N- and C-terminal affinity tags, so we introduced internal tags in the E2 flexible loops of H77 (at position 480 in variable region 2 (VR2), residues 459-483) and S52mod (at position 578 in variable region 3 (VR3), residues 570-585) (fig. S1C). This enabled affinity purification of strep-tagged H77 and S52mod E1/E2 complexes. To validate that the E1/E2 complex existed prior to detergent extraction, we performed strep-purification of E1/E2 extracted from cells transfected either with HIS or strep-tagged E1/E2 or co-transfected with both HIS- and strep-tagged E1/E2. Mixing HIS and strep-tagged E1/E2 membrane fractions prior to detergent extraction did not result in a HIS signal in BN PAGE WBs following strep purification, whereas doing the same for co-transfected cells gave a bright HIS signal (fig. S1D), indicating that the complex was formed before extraction and contained more than one E2 protein.

For cryo-EM studies, membrane-extracted H77 E1/E2 and S52mod E1/E2 were isolated by strep-tag affinity and ion-exchange chromatography. The purified complexes bound several NAbs targeting non-overlapping epitopes in E1/E2 as well as the CD81 large extracellular loop (Fig. 1B and fig. S1E). For S52mod-E1/E2, we were able to generate a cryo-EM structure determined at an overall resolution of 3.55 Å, which improved to 3.38 Å using local refinement centered on the ectodomains alone (fig. S2 and fig. S3). This revealed the presence of anti-parallel E1/E2 homodimers (Fig. 1C). Analysis of the structure showed high conservation of the predicted contact residues between the two E1/E2 heterodimers, with N-linked glycans exclusively found outside of this E1/E2 homodimer interface (Fig. 1D). Moreover, we obtained low-resolution data at 8.3 Å for H77-E1/E2 and the calculated map supports a similar anti-parallel fold to the S52mod-E1/E2 model (fig. S4).

The overall E1/E2 architecture agrees well with the recently solved E1/E2 heterodimer structure *(19)*, exhibiting identical disulfide bonding (fig. S5A). We also observe N-linked glycosylation at three of the same positions on E1 (N196, N209, N305) and seven of the same positions for E2 (N423, N448, N476, N557, N651, N701). Notably, N234 and N533 seem to lack glycosylation, while N430 and N581 are poorly resolved in our structure. Furthermore, the N-terminal domain (NTD) and C-terminal loop (CTR) of the E1 ectodomain exhibit nearly identical folds to the recently reported structure *(19)*. The overall fold of the E2 core domain also agrees well with E1/E2 heterodimer structure and with crystal structures of recombinant E2 *(4, 7-12)*, while variations are evident in the configurations of the E2 front layer (E2-FL, residues 420-460) and the CD81 binding loop (CD81bl, residues 519-535). In addition, our structure provides several novel structural insights such as the existence of the E1/E2 homodimer itself, and the shape of previously unresolved domains (Fig. 1, A and C), namely the TM domains formed by helices of E1 and E2, as well as the N-terminal segment of HVR1 and the E2 base and stem residues that make up the HVR1 binding cleft. Additionally, we observe a distinct arrangement of AS412.

### The homodimer interface

The homodimer interface consists of an E2 area (~1200 Å^2^), spanning the central ß-sandwich (residues 484-518 and 536-569), as well as the post VR3 (587-601), the E2 back layer (E2-BL, 602-649) and the regions known as the base (650-706) and the stem (707-720). As such, the homodimer exposes the immunodominant neutralizing face (residues 412–446 and 525–535), while the non-neutralizing face of E2 is buried in the homodimer interface. In line with this, we only detected binding of the non-neutralizing antibody, 2A12 (targeting a linear epitope in E2-BL), to soluble E2 and not the purified homodimer (fig. S6A and fig. S6B).

The stabilization of this E2-E2 interface relies on non-covalent interactions, involving residues that mediate hydrophobic interactions or hydrogen bonding (Fig. 2A). All residues involved exhibit a high degree of conservation across virus isolates within the same genotype as S52 (genotype 3) (Fig. 2B and fig. S6C). The E2-BL, the base and the stem region of the complex encompass two proximal hydrophobic pockets. One pocket involves residues L672, H673, S674, M688, L691, and L708 on each E2 monomer. The other pocket displays symmetry around H669, involving residues R636, F638, H669, P687, and P689. The E2-E2 interface is further fortified by interactions involving residues E597, T599, G640, F642, and R665 situated in post VR3, E2-BL and the base part of the complex, and P546, S547, and R602 in the ß-sandwich and E2-BL. Based on alanine scanning mutagenesis, we observed that most of these residues are important for maintaining homodimer integrity (Fig. 2C and fig. S6D). Notably, P546, T599, G640, R665, R636, L672, S674, L691, and L708, which are extensively conserved across all genotypes, were previously demonstrated to be critical for HCV pseudoparticle (HCVpp) infectivity *(28)*.

**Fig. 2.**
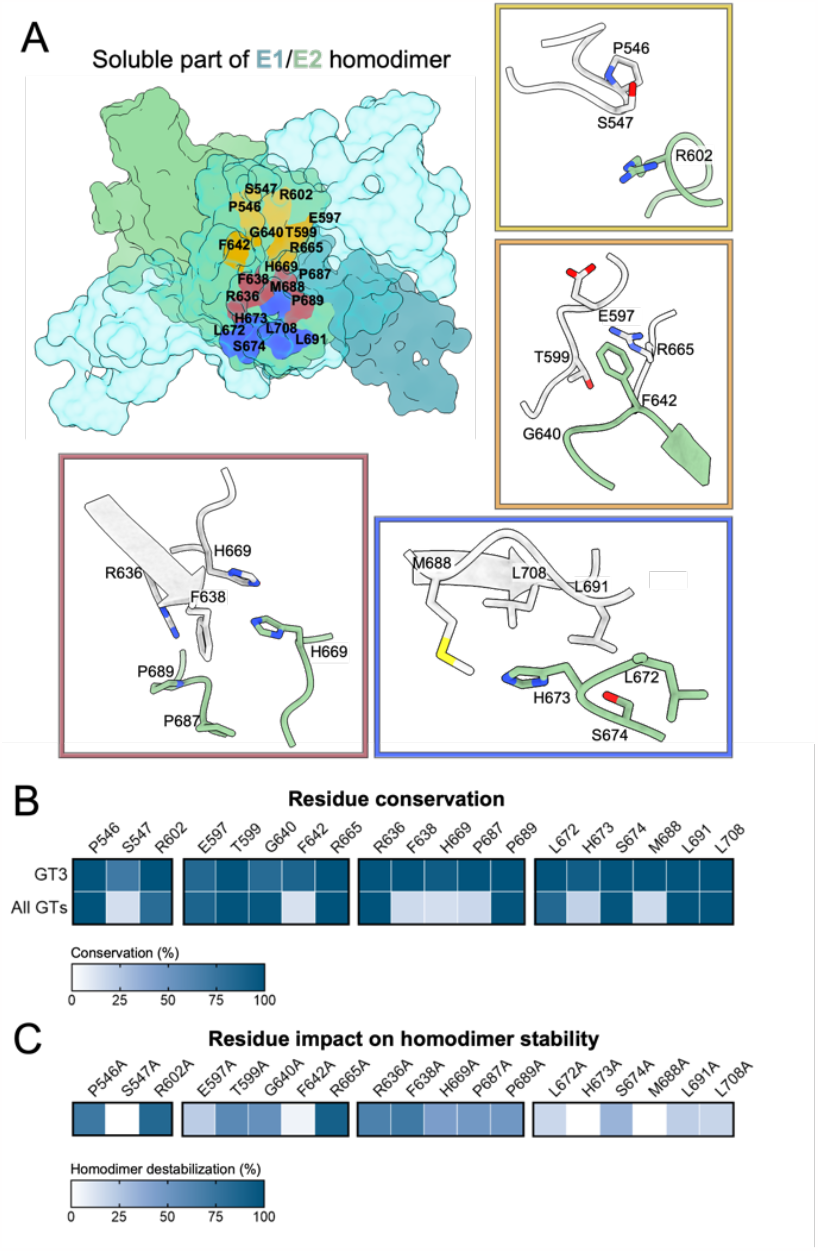
The homodimer interface of the HCV envelope structure. (**A**) Characterization of the E2 ectodomain interface of the homodimeric E1/E2 complex, with one E1/E2 monomer depicted in transparent turquoise and the other colored in steel blue (E1) and green (E2). The four panels provide details of the interactions (residues containing non-hydrogen atoms within 4 Å of each other) that we assert stabilize the interface. In each box, one monomer of the homodimer is depicted in green, while the other is in light grey. **(B)** Conservation of contact residues in genotype (GT) 3 and across GT 1-8 (Los Alamos HCV sequence database). **(C)** Homodimer destabilization by mutating contact residues to alanine. Measured as % band intensity compared to wildtype in native-page western blot (fig. S6D).

### The interaction of HVR1 and AS412 with other regions of E2

Detection of HVR1 and AS412 in E2 and E1/E2 structures has previously been challenging. However, the cryo-EM data of the homodimer reveals well-resolved density for HVR1 residues 384-393, as well as the complete AS412 (Fig. 3A). In our structure, the N-terminus of HVR1 extends into a cavity located between the base and stem domains of the same E2. This configuration is stabilized through a network of interactions with surrounding residues, including T631, L632, F633, W652, R654, G655, E656, N701, I702, V703, and D704. This finding aligns with our earlier discovery that the N-terminal part of HVR1 exerts a crucial influence on the compatibility of HVR1 with the remainder of E1/E2 *(29)*. For AS412, we observe a hairpin-like arrangement where the hairpin apex (at residue G418) protrudes apically from the E2 ectodomain. This resembles AS412 conformations observed when peptides spanning residues 411-423 are bound by the NAbs AP33 *(30)* and HCV1 *(31)*. The AS412 arrangement is stabilized by interactions with surrounding residues, including Q454, R455, G451, G560, F561, T599, and W622. The location of AS412 and the N-terminus of HVR1 corresponds closely with the Alphafold prediction for these segments (fig. S5B).

**Fig. 3.**
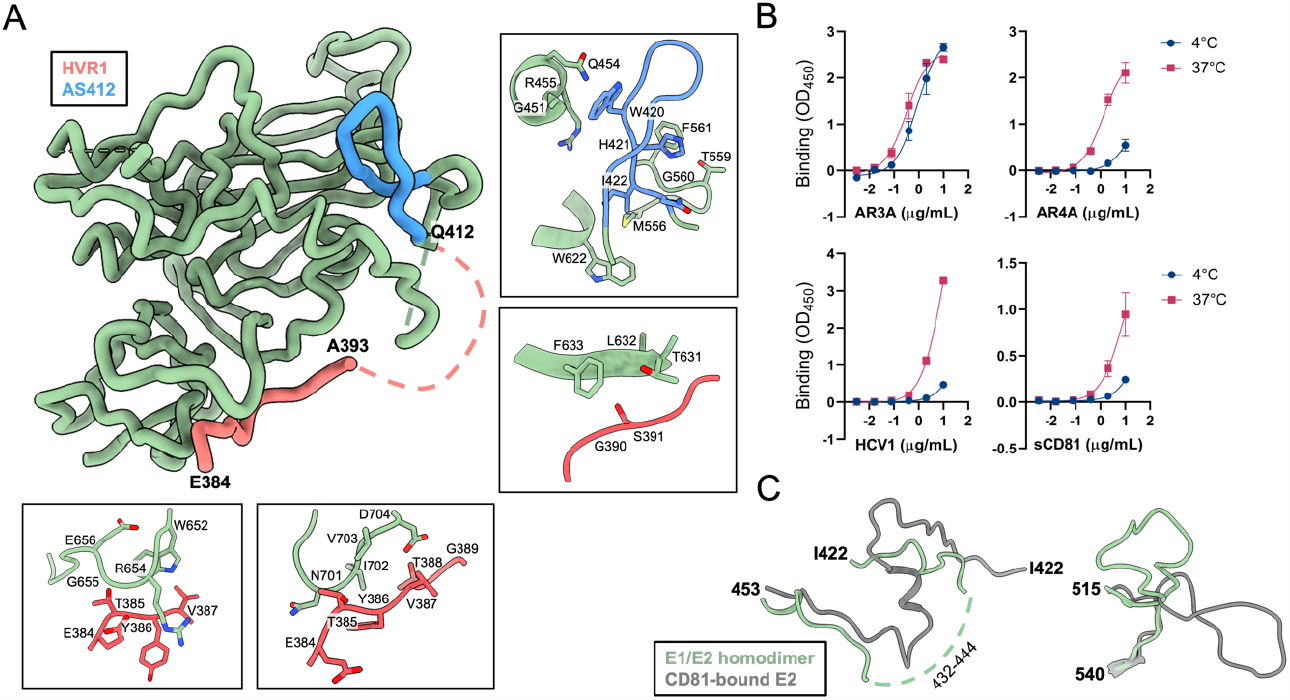
Interaction of HVR1 and AS412 with other regions of E2. (**A**) E2 ectodomain (green) with panels showcasing the interactions (residues containing non-hydrogen atoms within 4 Å of each other) that we assert stabilize the observed orientations of HVR1 (red) and AS412 (blue). (**B**) Binding of NAbs and sCD81 measured by ELISA performed at 4°C and 37°C. (**C**) A comparison between the aligned residues (422-460 and 515-540) of our E1/E2 homodimer model with the structure of E2 bound to CD81. Regions with missing structural data are indicated by broken lines.

Well-characterized conserved neutralizing epitopes on E2 exhibit diverse exposure in our structure (fig. S7). While antigenic region 3 is flexible and accessible for NAb binding, antigenic region 4 is concealed by HVR1 in the observed E2 configuration. Additionally, the interaction between the NAb HCV1 and its epitope, AS412, is hindered by the E2-FL. In accordance with this, we observe low binding of the NAbs AR4A and HCV1 to the purified dimer in ELISA assays conducted at 4°C; a temperature close to that of the sample prior to cryo-EM freezing (Fig. 3B). However, at 37°C we observe increased binding of both bNAbs suggesting that both HVR1 and AS412 dissociate to some extent from the rest of E2. For AR3A, we observe similar binding at 4°C as at 37°C, suggesting that accessibly of this epitope is not regulated by this mechanism.

Moreover, we observe low sCD81 binding of the purified dimer in 4°C ELISA (Fig. 3B). The binding site for the CD81 receptor, encompassing AS412, the E2-FL and the CD81 binding loop (CD81bl, residues 519-535), exhibit a distinct orientation in our structure compared to other E2 structures (Fig. 3C and fig. S5B) *(8)*. This discrepancy arises, in part, from the conformation adopted by residues 422-451 in those structures, which includes the AS434 helix (434-446) and the connecting loops, causing the CD81 binding residue I422 of AS412 to be in proximity to the CD81bl. In our structure, AS434 displays flexibility (residues 432-444 are poorly resolved), while the E2 FL residues 424-431 are rotated nearly 180° (Fig. 3C). This conformational flexibility corroborates with prior hydrogen-deuterium exchange experiments *(32)* and the demonstrated orientational rearrangement of AS434 upon binding to the NAb 212.1.1 *(9)*. Furthermore, in our structure, the CD81bl exhibits a bent configuration in the opposite direction relative to CD81, reminiscent of the recently resolved E2core+stem structure *(4)*, where this loop would require additional movement to initiate receptor interaction.

### The E1 and E2 membrane interaction

While the detergent reconstituted part of the complex was less well-resolved than other portions, a symmetrical set of TM bundles are clearly identified, encompassing an internal region in E1 (residues 262-289, overlapping the putative fusion peptide (pFP)-containing region (PCR)) and the C-terminal region in E1 (328-383) and in E2 (721-752) (Fig. 4 and fig. S2C). These sets of TM bundles, at either end of the homodimer, are tilted away from each other in a non-native orientation, likely because the two distinct sets of TM bundles of the complex share a single micelle, and the absence of peripheral constraints normally imposed by the lipid bilayer of the viral envelope (fig. S8A). This arrangement is likely enabled by flexible hinge regions between the TM bundles and the E1 and E2 ectodomains, the existence of which is supported by a comparison of our structure with the recently described structure of a single E1/E2 heterodimer (fig. S8B) *(19)*.

**Fig. 4.**
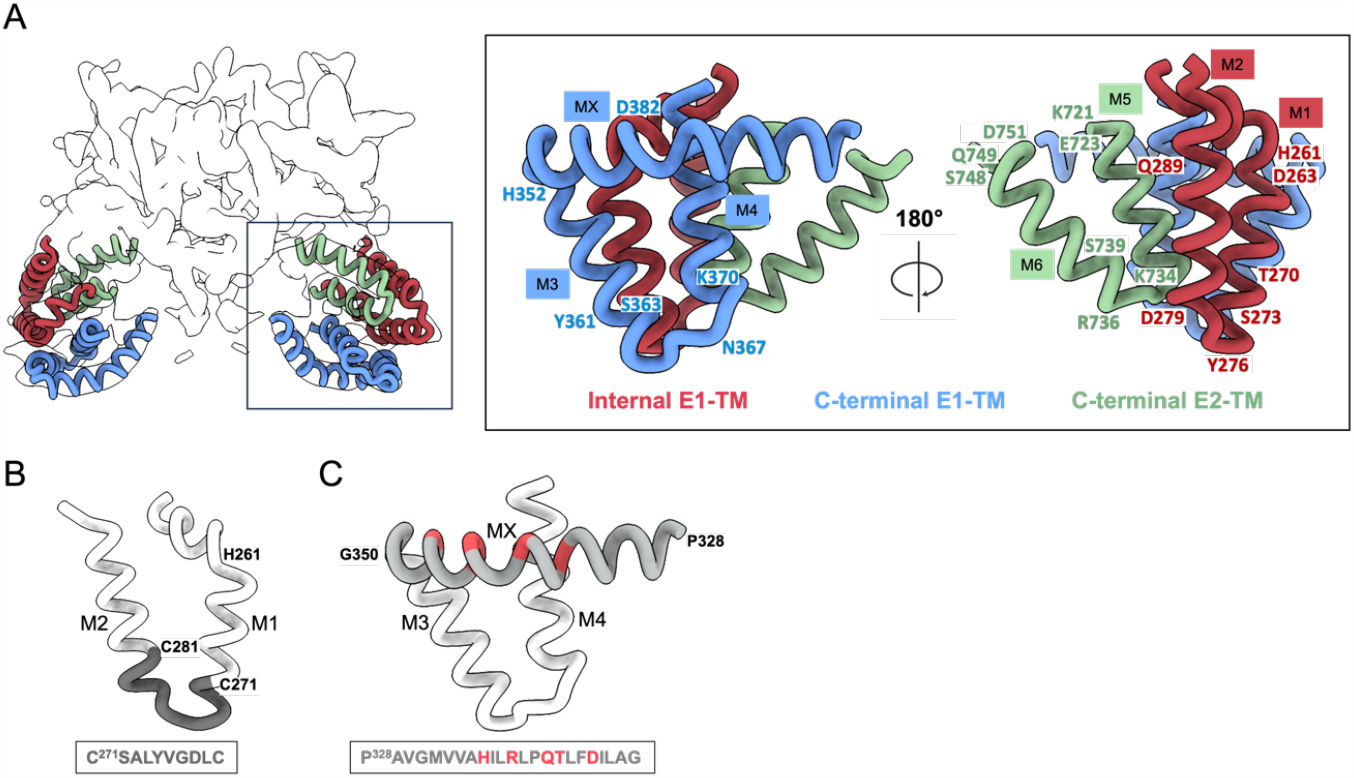
The E1 and E2 membrane interaction. (**A**) Model of the E1 and E2 TMs. Left: Cryo-EM map (transparent) overlayed with the predicted model of the different TMs of E1 and E2. Right: The modeled TMs oriented with hydrophilic residues (assigned with residue numbers) oriented in basal/apical positions. (**B**) The putative fusion peptide (depicted in dark grey) is located on the tip of the internal TM of E1 (M1+M2, depicted in light grey). (**C**) A longer helix (MX, depicted in grey) positioned perpendicular to the C-terminal TM of E1 (M3+M4, depicted in light grey). Hydrophilic residues are shown in red.

The two separate TM bundles are made up of pairs of short alpha helices (M1-M6; M1-M2 from the PCR, M3-M4 from the C-terminus of E1, and M5-M6 from the C-terminus of E2), predominantly composed of hydrophobic residues, with interconnecting linkers, containing several hydrophilic residues (Fig 4A). Additionally, M3 is preceded by a long helix (residues 328-350) that we designate MX. Also, M1 (residues 262-274) is part of a longer helix that kinks at the membrane interface at around H261. It is followed by a short loop (275-278) and M2 (279-289). Interestingly, M1-M2 covers the highly conserved hydrophobic sequence CSALYVGDMC (272– 281) (Fig. 4B). Several studies have shown the importance of this sequence in viral fusion with the host cell, and denoted it a putative fusion peptide, or pFP *(33, 34)*. M3 and M4 are two short helices, which are interconnected by a short loop, while MX is partly amphipathic (Fig. 4C), with the hydrophilic side pointing away from the TM helices, indicating that a portion of this helix lies in the membrane interface of the outer leaflet of the virus envelope. M5 and M6 are connected by a short loop and exhibit an inclined orientation with respect to the other TMs.

## Discussion

The structural elucidation of higher-order E1/E2 complexes on HCV particles has been hindered by the pleiomorphic nature of these particles *(35)*. In our model, the HCV E1/E2 complex forms a homodimer, which is not entirely unexpected, considering that numerous members of the *Flaviviridae* virus family feature envelope proteins adopting homodimeric configurations *(36)*. As there is little evidence supporting pH-driven or proteolytic maturation of HCV E1/E2 (like the structural changes observed for the envelope proteins of flaviviruses during their transit from the *trans*-Golgi network to the extracellular environment *(37)*), and our complex exhibits CD81 binding and comparable NAb epitope access regulation as observed for infectious HCV particles, it suggests that the ectodomain structure represents the mature, fusogenic state present on the virion. We demonstrate that the homodimer is stabilized through hydrophobic patches within the E2 subunits. This offers valuable insights for rational design of vaccine antigens that strategically conceal the regions of E2 sequestered within the dimeric assembly, thereby mitigating the induction of low potency or non-neutralizing Abs.

Inherent plasticity of HVR1 and the neutralizing face of E2 (encompassing AS412, AS434 and the CD81bl) have been established as key determinants of HCV NAb evasion, as well as influencing viral receptor engagement *(5)*. Indeed, we observe discrepancies with solved structures in the arrangement of these parts of E2. The HVR1 N-terminus, which is flexible in previously solved E2 structures, packs closely to base and stem domains of E2, concealing the conserved antigenic region 4. Furthermore, in this E2 configuration the conserved CD81 binding residue I422 in AS412 would need to undergo a significant movement of approximately 30 Å to facilitate CD81 binding *(11)*. Thus, we propose that the observed arrangement of HVR1 and the neutralizing face is associated with an overall NAb resistant state of the E1/E2 complex, less able to engage CD81. However, as we observe AR4A and CD81 binding to the complex at 37°C, but not at 4°C, we propose that the complex is capable of sampling configurations that are better poised for CD81 binding and likely also with increased sensitivity to NAb binding. In line with this, we have previously shown that neutralization of HCV particles by AR4A is increased at higher temperature, while this effect is abolished by HVR1 removal *(22)*.

Our structure indicates that the pFP of E1 is buried within the viral membrane, proximal to the C-terminal E1 and E2 TM helices. This carries significant implications for our understanding of the HCV fusion mechanism, as it suggests that the insertion of this domain may require considerable rearrangements, even within the E1/E2 heterodimers. The presence of a membrane-buried pFP is supported by its ability to anchor E1 in membranes in the absence of the C-terminal TM *(38)*, as well as reports that its helical TM hairpin structure appears to be conserved in recent AlphaFold predictions of E1 structures of *Hepaci-, Pegi-*, and *Pestiviruses (39)*. An intriguing alternative fusion mechanism was recently proposed to involve the insertion of two aromatic residues from the CD81bl into the host membrane at low pH *(4)*. The involvement of CD81bl in mediating membrane attachment in the late endosome is corroborated by the apical positioning of CD81bl in our structure. Moreover, the presence of flexible hinge regions that connect the TM helices with the ectodomain of E1/E2 would permit ectodomain elevation and dimer dissociation, which is known to be critical for fusion of *orthoflaviviruses* with known, class II, fusion mechanisms *(40)*.

In conclusion, we have successfully elucidated the structure of HCV E1/E2 in an anti-parallel homodimeric arrangement, and our findings form a basis for homodimer-based vaccine development targeting bNAb epitopes and avoiding the induction of non-neutralizing antibodies.

## Acknowledgments

We are grateful to Tillmann Hanns Pape at the Danish Cryo-EM Facility at the Core Facility for Integrated Microscopy (CFIM, University of Copenhagen) for assistance with sample screening and data collection; to Julian Conrad, Karin Wallden, Dustin Morado and Marta Carroni at the Cryo-EM Swedish National Facility in Stocholm for sample screening and data collection; to Lotte Mikkelsen, Anna Louise Sørensen and Louise Nielsen (CO-HEP) for general laboratory support; to Bjarne Ørskov Lindhardt (Hvidovre Hospital) and Charlotte Bonefeld (University of Copenhagen) for support of the project; and Thomas Krey (Hannover Medical School, Germany), Mansun Law (Scripps Research Institute, USA), Arwind Patel (University of Glasgow, UK) and Charlie Rice (Rockefeller University, USA) for providing reagents.

## Funding

NOVO Nordisk BRIDGE grant NNF20SA0064340 (EHA)

Novo-Nordisk Foundation grant no. NNF14CC0001 (CFIM)

Lundbeck Foundation Experiment grant R324-2019-1375 (JP)

Lundbeck Foundation Experiment (PG)

Lundbeck Foundation Fellowship (JP)

Lundbeck Foundation Fellowship (PG)

Lundbeck Ascending Investigator (PG)

Lundbeck Foundation Experiment (KT)

Lundbeck Foundation Postdoc (RVM)

Candys Foundation (EHA, JB, and JP)

Candys Foundation (CHO, JB, and JP)

Sapere Aude Advanced grant (JB)

Distinguished Investigator Grant from the Novo Nordisk Foundation (JB)

The Swedish Research Council grant (PG)

Knut and Alice Wallenberg Foundation Prolongation Fellow grant (PG)

Knut and Alice Wallenberg, Family Erling Persson and Kempe Foundations (SciLifeLab)

## Author contributions

EHA and JP optimized the sample preparation for cryo-EM analysis. EHA, CHO, JP and CG optimized the protein purification strategy. EHA and CHO carried out the protein purification, cloned the constructs and performed BN-page analysis. KW optimized cryoEM sample freezing strategy, collected and processed the cryo-EM data. KW and JP built and refined the atomic models into the cryo-EM maps. EHA, CG, JP, PG and KW conceived the protein purification strategy. EHA, JP, PG conceived the construct design. AS performed ELISA experiments. EHA, CHO, AS, RVM and MF performed protein production. JB provided materials and guidance. EHA and CHO wrote the paper and made the figures. All authors contributed to the manuscript text by assisting in writing or providing feedback. Supervision of the research was conducted by EHA, JP, CG, KW, and PG.

## Data and materials availability

Cryo-EM maps and atomic coordinates have been deposited to the Electron Microscopy Data Bank (EMDB) and the Protein Data Bank (PDB), under the following accession codes: EMD-XXX and EMD-XXX for the Cryo-EM maps, and XXX and XXX for the atomic coordinates.

