## Supplementary material for "Structure of the hepatitis C virus E1/E2 envelope proteins in a homodimeric complex"

**The PDF file includes:**

Materials and Methods
Figs. S1 to S8
Tables S1

### Materials and Methods

Generation of E1E2 constructs

Generation of expression constructs of genotype 1-6 using phCMV-E1E2 (amino acids 192-746, H77 abs. ref. AF009606) was produced as previously described (*41-43*). S52mod was generated by introducing residues from H77-TM into S52-E1E2 by replacing the amino acid 351-383 and 725-753 of S52 with H77. Twin-strep-tagged expression construct with the H77 and S52mod E1E2 were generated by introducing a twin-strep-tag (SAWSHPQFEKGGGSGGGSGGSAWSHPQFEK) or HIS6-tag (HHHHHH) at amino acid position 480 in H77-E1E2 and position 578 in S52mod-E1E2. Cloning of the construct was done using InFusion (Takara).

Monoclonal antibody production

The variable genes for the monoclonal antibodies (mAbs) used were cloned into the pVitro1-IgG1/κ vector (https://doi.org/10.1038/srep05885) and expressed using HEK293F cells (ThermoFisher). The cells were pelleted 72 hours post transfection and the supernatant containing the mAbs were collected, and then affinity purified using protein G columns on the ÄKTA Pure system before concentrating on Vivaspin columns (Satorius) and buffer exchanging into PBS using Zeba spin columns (ThermoFisher)

Extraction of E1/E2 for small-scale testing

For the transient transfection of HEK293T cells with the E1E2 plasmid, we seeded 800,000 cells per well in a 6-well plate and incubated them overnight at 37°C at 5% $CO_2$. The following day, we transfected the cells with various E1E2 plasmids using Lipofectamine 2000 (Invitrogen). Four hours post-transfection, the cell culture medium was replaced, and the cells were allowed to incubate for an additional 24 hours. At 24 hours post-transfection, we collected a few drops of the cells that was stained for E1/E2 expression using a specific primary antibody, AR4A, and a secondary anti-human Alexa Fluor 488 antibody (ThermoFisher). Following another 24-hour period post-transfection, we extracted the transfected cells using Native Page sample buffer (4x) (Invitrogen) and 1% dodecyl-b-D-maltoside (DDM). The cells were subsequently subjected to centrifugation at 20,000 g for 30 minutes at 4°C. This was followed by a 30-minute treatment with Benzonase Nuclease (Sigma-Aldrich) and another 30-minute centrifugation at 20,000 g rpm at 4°C.

Native page western blot

HEK293T cells transiently transfected with E1/E2 plasmid were analysed for E1/E2 content using Native Page Western blot. The samples, alongside a protein molecular marker (Invitrogen), were loaded on a 4-16% Native Page Bis-Tris gel (Invitrogen) and electrophoresed in a XCell II SureLock Mini-Cell (Invitrogen). Subsequently, the proteins were transferred onto a polyvinylidene difluoride (PVDF) membrane using electroblotting with 1x NuPAGE transfer buffer (Invitrogen). Following this, the membrane was blocked with BSK for 1 hour and the proteins were probed with specific primary antibodies, through an overnight incubation at 4°C. The membrane was washed with PBS-T and incubated for 1 hour at room temperature with a secondary horseradish peroxidase cross-absorbed antibody (F(ab')2-Goat anti-human IgG Fc, Invitrogen), with detection of proteins using enhanced chemiluminescence detection (SuperSignal West Femto Maximum Sensitivity Substrate, Pierce) performed via Image Lab 5.2.1 from Bio-Rad.

Extraction and purification of E1E2 for cryo-EM

HEK293F cells were transiently transfected with plasmid harboring strep-tagged E1E2. The cells were pelleted 72 hours post transfection, resuspended in lysis buffer (50mM Tris-Cl pH 7.5, 200mM NaCl, 10% glycerol, 1.0 mg/mL DNaseI, protease inhibitor cocktail) and mechanically disrupted using a Potter-Elvehjem PTFE homogenizer. Crude membranes were isolated by 30 min centrifugation at 163,268 g and solubilized in lysis buffer containing 1% DDM. Insolubilized material was removed from the lysate through another 30 min centrifugation at 163,268 g, before it was filtered and added to a Strep-Tactin XT 4Flow gravity column (IBA). The column was washed with 5CV buffer W (100mM Tris-Cl pH 8.4, 150 mM NaCl, 10% glycerol, 1 mM EDTA) + 0.03% DDM, 5CV buffer W + 0.3% DDM, 5CV buffer W +0.03% DDM and 10CV buffer W + 0.05% LMNG. Protein was eluted from the column in BXT buffer (100mM Tris-Cl pH 8.4, 150 mM NaCl, 10% glycerol, 1 mM EDTA, 50mM biotin) + 0.01% LMNG, and concentrated using Vivaspin 20 Centrifugal Concentrator with 100 kDa MW cut-off. The concentrated protein was diluted 1:8 in IEX loading buffer (50mM Tris pH 6.5, 10% glycerol), and floated over CaptoQ XP resin (Cytiva) pre-equilibrated with IEX buffer (50mM Tris pH 6.9, 18.8mM NaCl, 10% glycerol). The IEX flow through containing E1E2 complex was collected and concentrated to 3 mg/mL, and buffer exchanged with Zeba micro desalting spin column into a cryo buffer (20mM Tris-Cl pH 7.5, 150mM NaCl, 1% glycerol, 0.01% LMNG).

ELISA

Purified E1/E2 (25 µg/ml in homodimer buffer (HDB, Tris-Cl pH 7.5, 150mM NaCl, 10% glycerol, 0.01% LMNG) was coated on MaxiSorp (ThermoFisher) plates overnight at 4°C. Plates were blocked for 1h at RT in blocking buffer (BB, 2% skimmed milk powders in HDB), washed four times with HDB, and subsequently incubated with serially diluted mAbs and sCD81 (CD81-LEL-Fc) in BB for 1h at 37°C, RT or 4°C. The plates were then washed six times with HDB, followed by the addition of HRP-labelled mouse anti-human IgG (Invitrogen) in BB. Subsequently, after an additional six washes with HDB, plates were developed by adding TMB (VWR). The reaction was stopped by adding 2M $H_3PO_4$. Absorbance was then measured at 450 nm, and the data were visualized using GraphPad Prism.

Cryo-EM sample preparation

3 µL of purified S52mod E1/E2 sample 1 mg/ml were applied onto a holey carbon grid (Quantifoil Cu R1.2/1.3, 300 mesh) glow-discharged for 60 s at 10 mA (EM ACE200, Leica Microsystems). Grids were blotted for 3.5 s and plunge-frozen in liquid ethane using a Vitrobot Mark IV (FEI) operated at 4°C and 100 % humidity. Due to more impurities in the H77 sample, we used a higher sample concentration of 2 mg/ml.

Data collection and image processing

S52mod data was collected on a 300 kV Titan Krios electron microscope (FEI) equipped with Gatan K3 direct electron detector. Energy filter was set to 20 eV width. For the S52mod dataset, a total of 13,555 movies were recorded using a pixel size of 0.824 Å, total dose of ~50 $e^-/Å^2$ fractionated into 40 frames, and with a defocus range of -2.5 to -0.5 µm. Data processing were done using cryoSPARC v3.3.1.

The movies were initially processed using Cryosparc patch motion correction and patch CTF estimation. A total of 2,372,136 particles were bin2 extracted to a box size of 160 pixels using template-free picking. The initial dataset was briefly cleaned up with 2D classification, and 1,642,301 cleaned particles were local motion corrected to a box size of 320 pixels for further data processing. After multiple rounds of 2D/3D classification, remaining particles were classified into 4 classes using *ab-initio* reconstruction followed by heterogeneous refinement. The electron density map was further improved by heterogeneous refinement, NU-refinement, applying C2 symmetry and local refinement employing a tight mask covering the protein region only. The final volume map at an overall resolution of 3.55 Å was calculated from 105,033 particles. The map resolution could be further improved to 3.38 Å resolution by local refinement within a tight mask that only cover the ectodomain. This map was used for model building, but not used for final refinement of the final model. The data processing pipeline is shown in Fig. S2. As indicated by the cryo-EM maps and local resolution estimates, the ectodomains were well-defined, while the membrane spanning portions were less well-resolved (fig. S2C).

The H77 dataset was collected and processed similarly as the S52mod dataset. Due to the low sample concentration and poor sample stability, only a total of 12,286 movies were recovered yielding a final resolution map of 8.3 Å, calculated based on 28,281 particles (from an initial 430,579 particles).

<u>Model building</u>

The initial model of the soluble part of S52mod E1/E2 was generated by rigid body fitting of a previously available structure (pdb ID: 7T6X) (*19*). All residues were manually mutated to the correct sequence in S52, and all glycosylations were kept until the end of visible density in the map.

The resolution of the membrane spanning part was insufficient for *de novo* model building. Hence, the membrane spanning part of the model was initially obtained from Alphafold (*44*). Next, the predicted membrane helical bundles were fitted into the low-resolution densities associated with the membrane helices, manually adjusted, and re-connected through MD simulation assisted model building.

The model building was done iteratively using coot (*45*) and phenix_real_space_refine (*46*). Secondary structure restraints and Ramachandran restraints were imposed during refinement. To assist model interpretation, the map was subjected to EMready for model building (*47*). Nonetheless, the original unsharpened map was used for model refinement. A structure of the ectodomain only, lacking residues 255-293 and from 316 in E1, and without amino acids 711 to the end of E2, was also generated using the model with the transmembrane part as a starting template and refined using similar principles and by using the 3.38 map (fig. S2). The quality of the models were validated and assessed using MolProbity (*48*) (see Table S1 for statistics). All figures were prepared with PyMOL (*49*), Chimera (*50*) and ChimeraX (*51*).

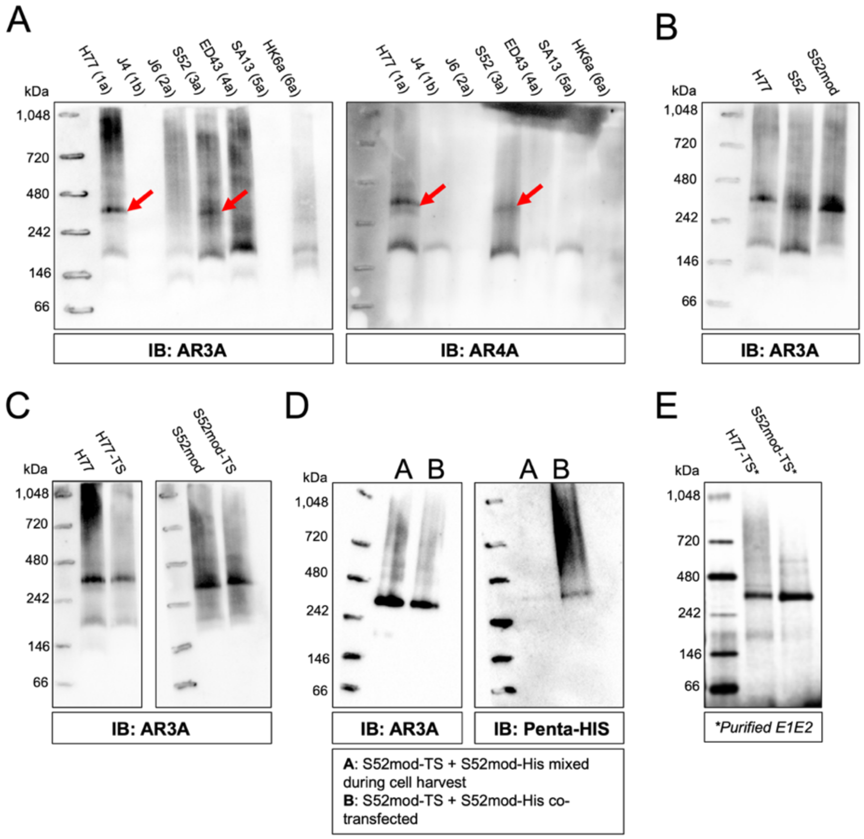


**Fig. S1. Purification of HCV E1/E2 in a higher-order complex.** (**A-E**) Extraction and solubilization of HCV E1/E2 analyzed by BN PAGE western blots. **(A)** Extracted HCV E1/E2 of different isolates of genotype 1-6. Red arrows indicate the higher-ordered E1/E2 complex. (**B**) Extracted E1/E2 from HCV isolate S52 (genotype 3a) with TMs from H77 (genotype 1a) (termed S52mod). (**C**) Extracted E1E2 from H77 and S52mod with internal twinstrep tags (termed H77-TS and S52mod-TS). (**D**) Strep-purification of extracted E1/E2 from HIS- and strep-tagged S52mod E1/E2 expression. (**E**) Coomassie stain of purified H77-TS and S52mod-TS E1/E2 complex.

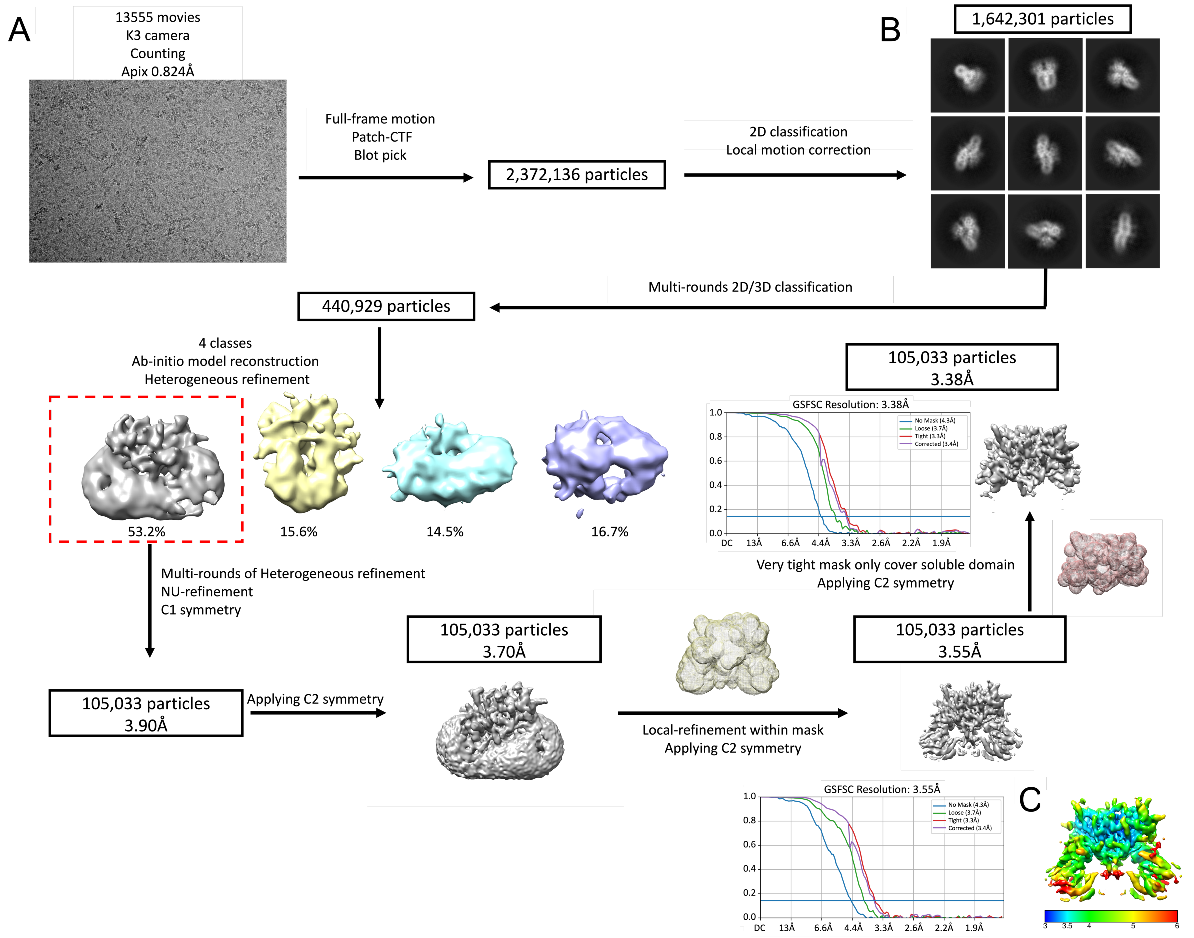


**Fig. S2. Cryo-EM data processing scheme of the E1/E2 in a homodimeric complex for S52mod. (A)** Representative cryo-EM image of S52mod. **(B)** Selected 2D averages of the particles. **(C)** Density of the final map used for model building colored by resolution. Table S1 contains specific information and parameters regarding the data collection.

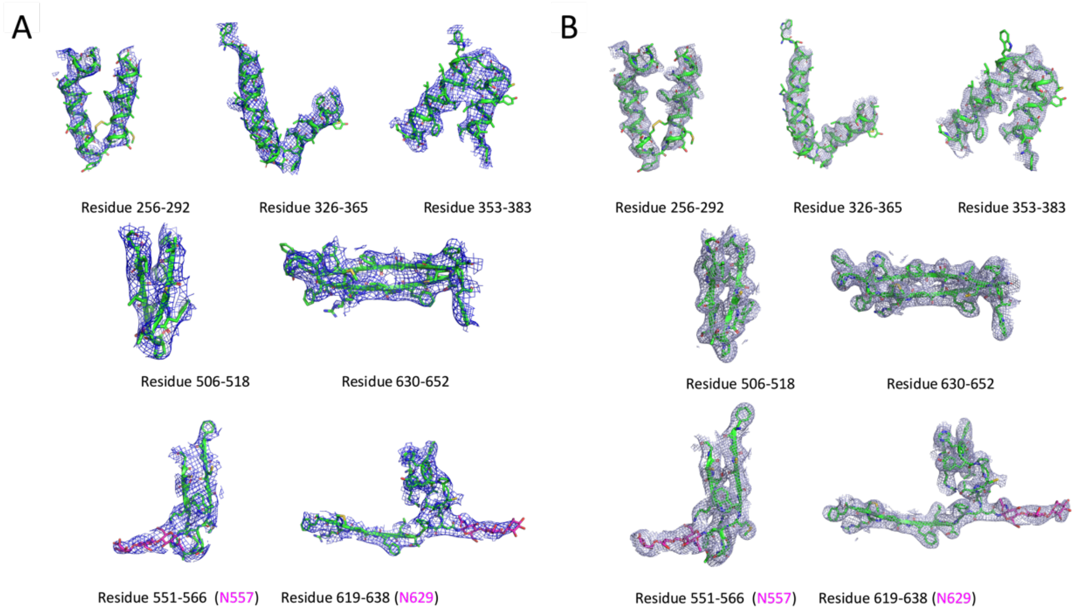


**Fig. S3. Structure and density fit.** Selected regions are shown with the density around the final structure. **(A)** Model fit into the original map without sharpening. **(B)** Model fit into the EMready refined map. Main chains are shown in cartoon and side chain in sticks. The glycosylation is shown as sticks colored in magenta. Glycosylated residues are labeled in magenta.

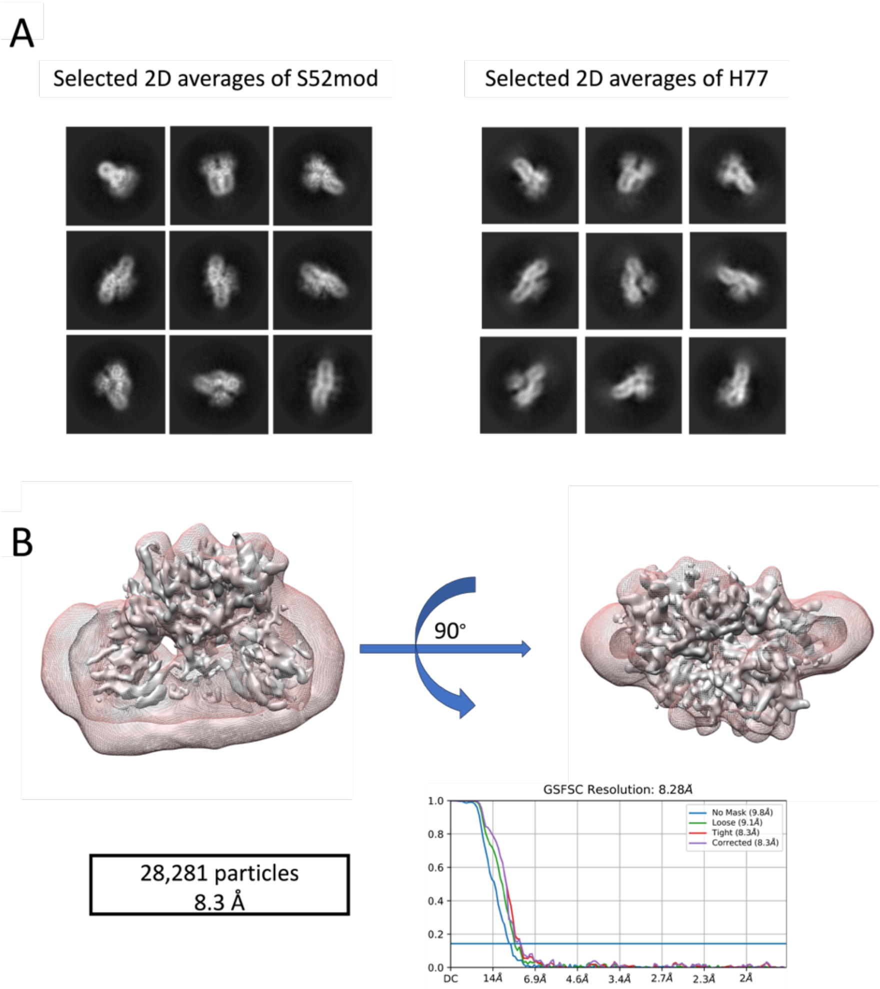


**Fig. S4. Structure comparison of H77 and S52mod.** The final map of H77 could only reach 8.3 Å resolution likely because of low sample stability and limited particle numbers, but both the 2D averages **(A)** and the overlay of the low-resolution map **(B)** of H77 (shown as red mesh) fit well with the high-resolution map of S52mod (shown in gray surface) with almost identical arrangement.

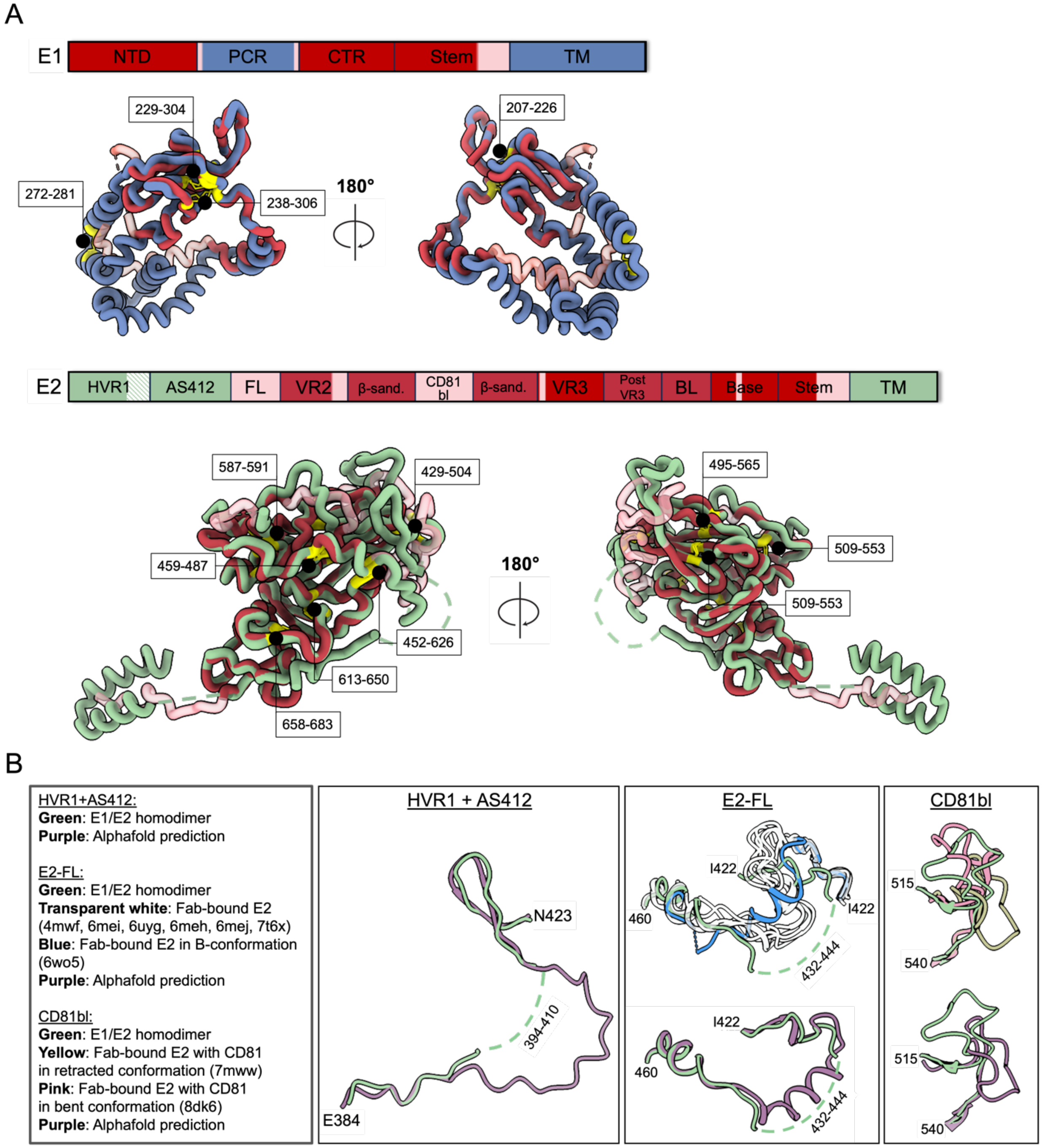


**Fig. S5. E1 and E2 alignment. (A)** Alignment of E1 and E2 from our modeled structure (depicted in blue (E1) and green (E2)) with the recently solved E1/E2 structure (7t6x; overlapping regions depicted in red and non-overlapping regions in transparent red). The location of disulfide bonds are numbered and highlighted in yellow. **(B)** A comparison between the aligned residues (HVR1+AS412: 384-423, E2-FL: 422-460, CD81bl: 515-540) of our modeled E2 structure, the AlphaFold predicted structure, and structures elucidated through Fab-based methods. Unresolved regions are indicated by broken lines.

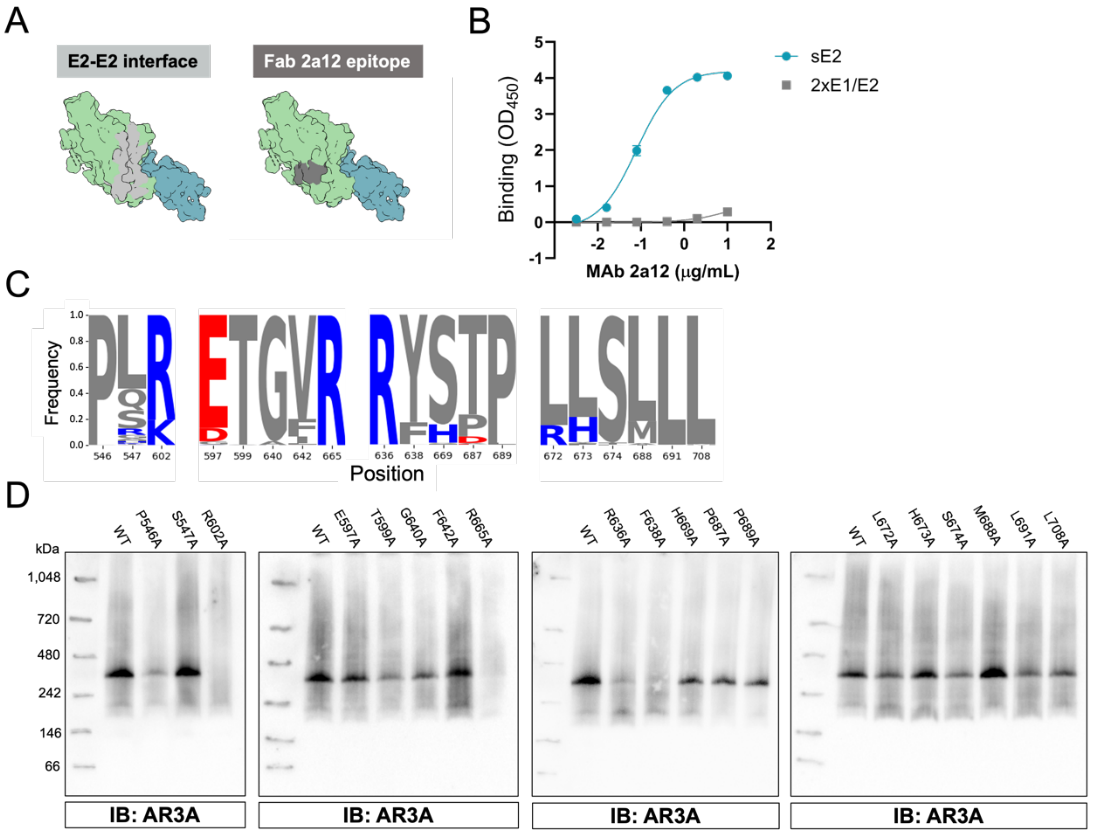


**Fig. S6. Analysis of the E1/E2 homodimer interface.** (**A**) Buried surface area of the E2-E2 interface of the homodimeric E1/E2 complex and the Fab2A12 epitope (*7*). The same view of the soluble part of the E1/E2 heterodimer is shown with E1 in steel grey and E2 in green. (**B**) Binding of the non-neutralizing antibody 2A12 to purified E1/E2 homodimer complex and sE2 measured by ELISA. (**C**) Sequence logo representing amino acid conservation within GT 1-8 for each contact residue (Los Alamos HCV sequence database). (**D**) Extracted HCV E1/E2 (S52mod) with alanine mutations in the contact residues of the homodimer interface, analyzed by BN PAGE western blot.

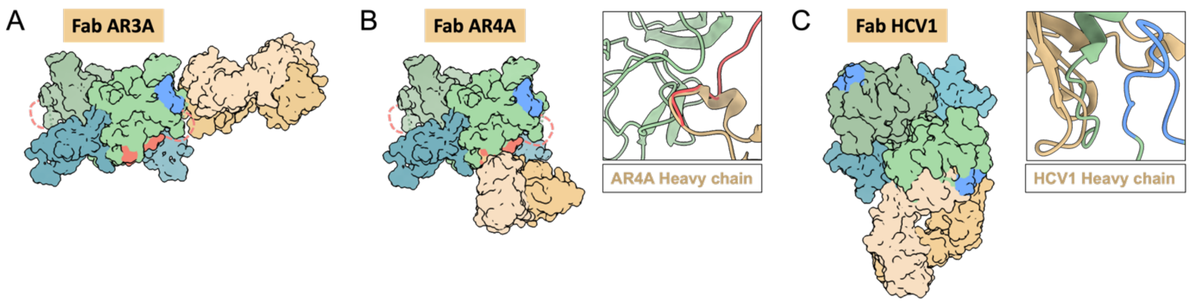


**Fig. S7. Characterization of the epitopes of NAbs AR3A, AR4A and HCV1 on the E1/E2 homodimeric complex.** (**A-C**) Structure of Fabs bound to E2 (A: 6bkb, B: 7t6x, C: 4dgv) superimposed on the E1/E2 homodimer structure. E1 is depicted in steel grey, E2 in green, HVR1 in red and AS412 in blue. In panel B, only the variable domain of the light chain and the heavy chain is displayed. Zoom-in on the clash of the AR4A heavy chain with HVR1 and the HCV1 heavy chain with E2-FL, are shown in B and C, respectively.

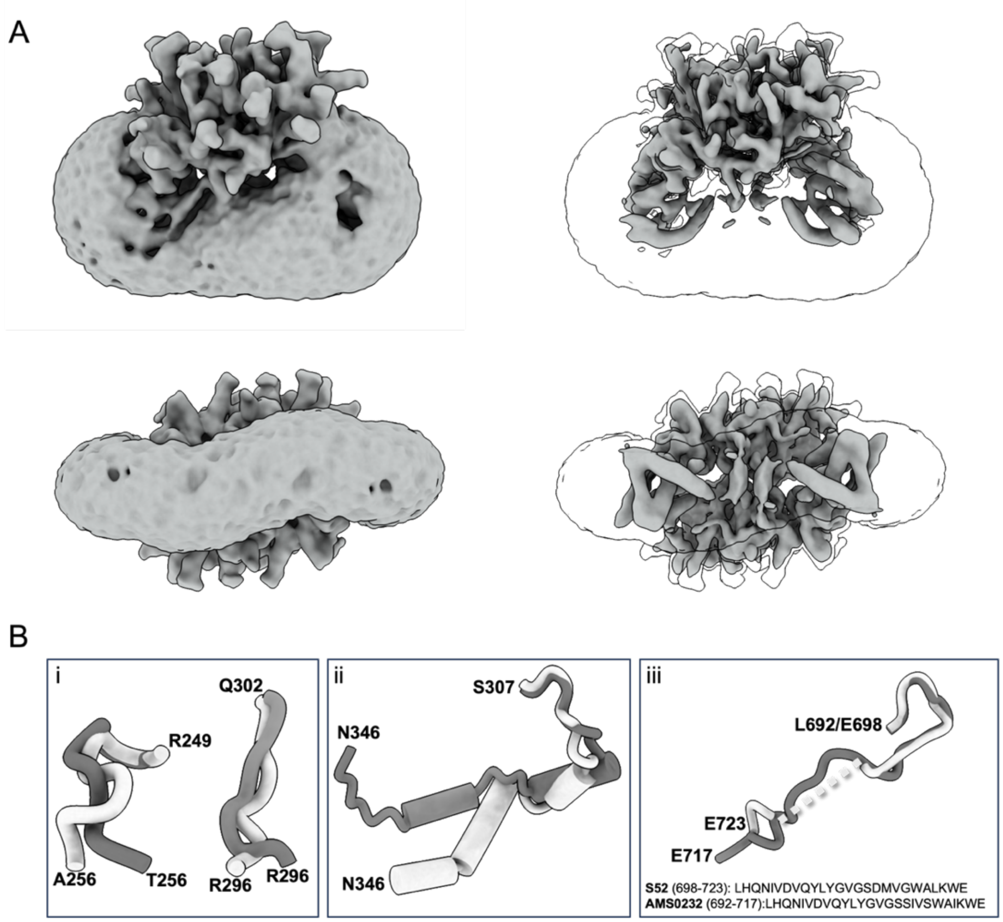


**Fig. S8. Analysis of the TM domains. (A)** The TM helices in the detergent micelle. Upper row: side view, lower row: bottom view. **(B)** Comparison of flexible hinge regions, connecting the ectodomains to the TM helices, of our modeled structure (depicted in light grey) with the recently solved E1/E2 structure (7t6x; depicted in dark grey).

### Table S1. Cryo-EM data collection and refinement statistics

| | **S52mod** | | **H77** |
|---|---|---|---|
| | **E1/E2 w. TM domains** | **E1/E2 ectodomain** | |
| | EMD-XXX, PDB XXX | EMD-XXX, PDB XXX | |
| **Data collection** | | | |
| EM equipment | FEI Titan Krios | FEI Titan Krios | FEI Titan Krios |
| Voltage (kV) | 300 | 300 | 300 |
| Detector | Gatan K3 | Gatan K3 | Gatan K3 |
| Data collection mode | counting | counting | counting |
| Pixel size (Å) | 0.824 | 0.824 | 0.8566 |
| Energy filter | 20 eV | 20 eV | 20 eV |
| Electron dose ($e^-$/Å$^2$) | 50 | 50 | 50 |
| Defocus range (μm) | -1.0 ~ -2.6 | -1.0 ~ -2.6 | -1.0 ~ -2.6 |
| Number of total movies | 13,555 | 13,555 | 12,286 |
| **Data processing** | | | |
| Software | Cryosparc | Cryosparc | Cryosparc |
| Number of initial extracted | 2,372,136 | 2,372,136 | 430,579 |
| Number of final used | 105,033 | 105,033 | 28,281 |
| Symmetry | C2 | C2 | C2 |
| Map resolution (Å) | 3.55 | 3.38 | 8.28 |
| **Model refinement statistics** | | | |
| Total built residues | 1098 | 812 | |
| Model-map-fit CC | 0.82 | 0.81 | |
| **R.m.s.d.** | | | |
| bonds (Å) | 0.008 | 0.003 | |
| angles (°) | 0.952 | 0.755 | |
| **Molprobity score** | 3.53 | 2.33 | |
| **Ramachandran plot** | | | |
| Disallowed (%) | 0 | 0 | |
| Allowed (%) | 16.58 | 12.81 | |
| Favored (%) | 83.42 | 87.19 | |
| Clash score | 32.45 | 16.17 | |
| Average B-factor (Å$^2$) | 220.61 | 194.31 | |
